# Quantitative Behavioural Phenotyping to Investigate Anaesthesia-Induced Neurobehavioural Impairment

**DOI:** 10.1101/2020.10.28.358978

**Authors:** Pratheeban Nambyiah, Andre EX Brown

**Affiliations:** Institute of Clinical Sciences, Faculty of Medicine, Imperial College London, London, UK; MRC London Institute of Medical Sciences, London, UK

**Keywords:** Pediatric Anesthesia, Neurodegeneration, C. elegans, Phenotyping

## Abstract

**Background:** Vertebrate animal experiments have suggested that anaesthesia exposure to the developing nervous system causes neuroapoptosis and behavioural impairment. Mechanistic understanding is limited, and target-based approaches are challenging. High-throughput methods may be an important parallel approach to drug-discovery and mechanistic research. The nematode worm *Caenorhabditis elegans* is an ideal candidate as a model for this. A rich subset of its behaviour can be studied, and hundreds of morphological and behavioural features can be quantified, then aggregated to yield a ‘signature’. Perturbation of this behavioural signature may provide a tool that can be used to quantify the effects of anaesthetic regimes, and act as an outcome marker for drug screening and molecular target research.

**Methods:** Larval *C. elegans* were exposed to: isoflurane, ketamine, morphine, dexmedetomidine, and lithium (and combinations). Behaviour was recorded, and videos analysed with automated algorithms to extract behavioural and morphological features.

**Results:** Anaesthetic exposure during early development leads to persisting morphological and behavioural variation (in total, 125 features/exposure combinations). Higher concentrations, and combinations of isoflurane with ketamine, lead to more feature ‘hits’. Morphine and dexmedetomidine do not appear to lead to behavioural impairment. Lithium rescues the neurotoxic phenotype produced by isoflurane.

**Conclusions:** Findings correlate well with vertebrate research: impairment is dependent on agent, is concentration-specific, is more likely with combination therapies, and can potentially be rescued by lithium. These results suggest that *C. elegans* may be an appropriate model with which to pursue phenotypic screens for drugs that mitigate the neurobehavioural impairment. Some possibilities are suggested for how high-throughput platforms might be organised in service of this field.

## Introduction

It is increasingly apparent that general anaesthetic agents can affect neurodevelopmental processes other than consciousness, and that these effects can persist beyond the period of clinical anaesthesia^1^. In fact, if we consider their presumptive mechanisms of action, it seems optimistic that we ever thought anaesthetics should have only short-term and reversible actions on consciousness. They appear to act promiscuously on receptor targets such as γ-Aminobutyric acid (GABA), N-methyl-D-aspartate (NMDA), and a variety of other protein channels, and are typically given at large concentrations to overcome their lack of selectivity^2^. These targets are widespread, and have defining roles to play in neural development and plasticity^3–5^. It is unsurprising therefore that interruption of consciousness at particular timepoints in brain age may be accompanied by extra-modal effects on the neuronal architecture, network or function.

Although there is genuine potential to harness these effects for good, (e.g. the modulation of glutamergic signalling by ketamine in the treatment of depression^6^), there is also significant public health concern^7^. Research in animals has demonstrated apoptotic neurodegeneration in the developing brain and persistent impairments in learning and memory, associated with exposure to volatile anaesthetics, ketamine, nitrous oxide, midazolam and propofol^8–10^. In general, longer exposures and combinations of anaesthetics delivered together have resulted in the most striking phenotypes. This phenomenon has been termed anaesthesia-induced neurotoxicity (AIN), and the findings have been replicated in multiple different organisms, including non-human primates^11^. Ethical and practical constraints mean that trials in human children have found it difficult to separate the role of anaesthesia from those of surgery, co-morbidity, socio-economic factors and other confounders^12^. Thus a mixed picture has emerged from clinical studies^13–17^. Animal models then, have an important role to play, particularly in elucidating mechanism, developmental windows of vulnerability, effects on neural architecture, and ability to quantify and therapeutically modulate the response.

This last consideration is the motivation for this work: how to apply rigorous quantitative analysis to record neurodevelopmental outcomes after exposure to anaesthesia, in a way which allows rapid and scalable screening of agents and modulating factors? Although mammalian models have provided much insight, there are drawbacks: vertebrates exhibit many degrees of freedom in their behaviour but only a limited number of heterogeneous outcomes have been used to quantify responses to anaesthesia exposure, leading to constrained expressions of nervous system output.

Different groups have used, variously, open field tests^18^, fear conditioning tests^19^, assorted tests of spatial awareness, learning and memory^20,21^, and tests which were initially developed to assess depressant behaviour such as the forced swimming test^22^. The lack of standardization limits the power of both individual studies and meta-analyses, and in the absence of a generally accepted behavioural phenotype for AIN, it is difficult to be sure whether the constrained set of behaviours exhibited in these experiments can truly reflect the neuropathological deficit.

For these reasons, until there is better mechanistic understanding of AIN, a high-throughput phenotypic screening method may be an important parallel line of investigation. This approach is difficult in mammals, but has been successfully demonstrated with other whole-animal models, in particularly the nematode worm *Caenorhabditis elegans*^23,24^. Because of its small size, short life cycle (Fig 1a), and ease of maintenance and propagation, *C. elegans* can be raised in large numbers in the lab. Key anaesthetic targets including GABA and NMDA receptors are conserved in *C. elegans* and neuronal cells in the worm undergo apoptosis through conserved mechanisms first discovered in *C. elegans*^25^. Worms are susceptible to anaesthesia and anaesthesia-induced neurotoxicity^26^. However, the potential of *C. elegans* goes further: its behaviours are amenable to quantitative aggregation and analysis, which yields a phenotypic ‘signature’. Perturbation of this behavioural signature in response to anaesthesia exposure may provide a powerful tool that can be used to quantify the effects of varied anaesthetic regimes, and act as an outcome marker for drug screening.

**Fig 1.**
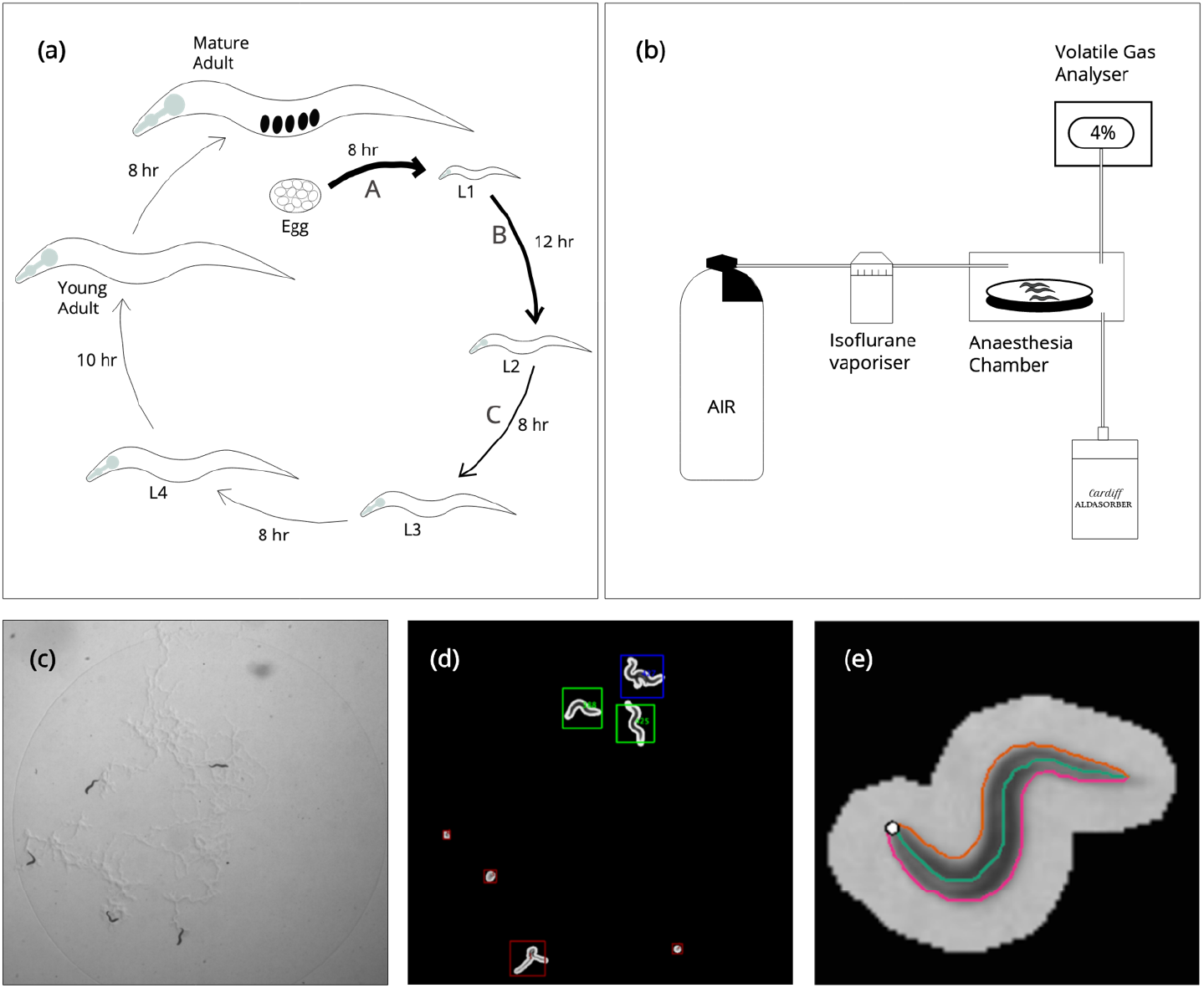
Experimental setup and worm tracking. (a) Life cycle of C. elegans hermaphrodite at 22°C from egg to mature adult via larval stages. Thickness of arrows represent degree of neurogenesis (A, during proliferation phase of embryogenesis leading to 222 neurons at hatching; B, 74 neurons added during L1; C, 6 neurons added during L2). (b) Anaesthesia apparatus. Air flows from a pressurised cylinder into a vapouriser, and is saturated with a variable concentration of the volatile gas anaesthetic isoflurane. It then passes into a chamber holding plates of C. elegans. The delivered concentration is measured directly with an optical analyser, and titrated to desired value. The output from the chamber flows into the Cardiff Aldasorber, which binds anaesthetic gases for safe disposal. (c) Video still of experimental plate, showing 5 L1 larvae on an E. coli OP50 lawn. (d)-(e) Larva segmented from background using intensity threshold to extract the worm contour and skeleton.

There are numerous theories linking anaesthesia exposure to neurodegeneration and behavioural impairment^27,28^. These include: disruptions in neuronal activity, signalling and neurotransmitter function; impaired neuronal maturation, integration, dendritic spine and synapse formation; abnormalities in growth and nutrient signalling pathways; microglia-related mechanisms; oxidative stress; impairment of autophagy; and epigenetic regulation. No single theory unifies these mechanisms, or accounts for all reported histopathological and neurobehavioral phenotypes. Given the range of possible mechanisms as well as the uncertainty around each of them, a phenotypic screening approach can generate both candidate drugs for further testing as well as new mechanistic hypotheses. In this work, we generated a behavioural dataset resulting from the exposure of a developing nervous system to anaesthesia, using a whole animal *C. elegans* model. We provide proof of concept that a *C*. elegans-based high-throughput phenotypic screening system can aid in drug discovery and mechanistic research in AIN.

## Methods

The N2 strain of *C. elegans* was used in all experiments. Worms were maintained as per previously published protocols^29,30^. Larvae were synchronised to a 1-hour time window as follows. 20 well-fed adult worms were left to lay eggs overnight. In the morning, all worms were washed off the plate using M9 buffer, leaving only eggs. Plates were left for 1 hour, and then inspected. Any larvae that had hatched in this 1-hour period were retrieved by washing the plate three times with 0.5ml M9 buffer and centrifuging the larvae-containing liquid at 2,500 rpm for 30 seconds. The supernatant was discarded, and the larvae transferred to a new low-peptone plate pre-seeded with 10μl *E*. coli OP50 bacteria. Typically, this step would yield several dozen animals. At all times other than during video recording, plates were kept in a 22°C incubator. At 1 hour after hatching, larvae were exposed to anaesthesia. This timepoint was chosen because it is a period of extensive neurogenesis and neural rewiring (Fig 1a); exposure to isoflurane at this stage has previously been showed to result in defective behaviour^26^. Agents were chosen because of previous vertebrate literature suggesting a neurotoxic effect (ketamine^8^, isoflurane^9^), lack of effect (morphine^31^, dexmedetomidine^32^) or potential to rescue the neurotoxic deficit (lithium^33^).

### Anaesthetic exposure & video recordings of behaviour

#### Isoflurane

Larvae were exposed at concentrations of 0% (control), 1%, 2% or 4% in air (flow rate 0.3L/min). The apparatus for delivering isoflurane is shown in Fig 1b. Exposure was for 2 hours, and animals were then transferred to new plates to emerge from anaesthesia. Control animals were exposed to air-flow only within the anaesthesia chamber.

Videos of larval behaviour (see Fig 1c for video still) were recorded for 15 mins at the following intervals after hatching: (i) 8 hours, corresponding to late L1 stage; (ii) 23 hours, corresponding to mid-L3; (iii) 3 hours after eggs seen on control plate (corresponding to adult). (Isoflurane is expected to have cleared fully in <1 hour). 4-6 larvae were recorded per plate, and 3 plates were exposed at each concentration. The videos were recorded using a custom-made multi-worm tracker built by Real Time Computing (East Sussex, UK).

#### Other agents

Synchronised larvae were transferred onto a plate containing 1μM, 10μM or 100μM of the following drugs: ketamine hydrochloride, morphine sulphate, dexmedetomidine hydrochloride (1μM and 10μM only – attempts to use 100 μM concentrations resulted in drug precipitation), and lithium chloride. The following combinations were also studied: (a) Isoflurane 1% with Ketamine 1μM, 10μM and 100μM; (b) Isoflurane 2% with Ketamine 1μM, 10μM and 100μM; (c) Isoflurane 4% with Ketamine 1μM, 10μM and 100μM; (d) Isoflurane 4% with Lithium 1μM, 10μM and 100μM. Exposure was for 2 hours and animals were then transferred to new plates. Postexposure recording was as given above.

### Data analysis

Worms were tracked using the open source Tierpsy Tracker^34^. Briefly, worm pixels were segmented from the background using an intensity threshold, and the contour of each segmented object is extracted and given a unique identifier (Fig 1d). The worm skeleton is then defined as the midline connecting the two points of highest curvature (head and tail, Fig 1e). The body of the worm is split into head, neck, midbody, hips and tail (and sub-divided further). Bend angles can be calculated as the difference in tangent angles at any given point. Combining this information with knowledge of time and the worm’s location on the plate allows extraction of a range of behavioural features, including parameters related to morphology, posture, locomotion and path dynamics^35^. The full set runs to over a thousand features; however, a subset of 244 features were accepted, as analysis has previously shown that a similar subset provides an optimal balance between accurate classification and reducing the handicap associated with correction for multiplicity^35^. These are given in supplementary data Table S1. Data was filtered so that only worm identities which existed for a minimum of 30 seconds were accepted, to minimize the effect of artefacts and segmentation errors which occasionally give rise to short-lived non-worm identifiers. Unpaired Student’s t-test was used to compare feature means between groups and we controlled the false discovery rate using the Benjamini-Yekutieli procedure^36^.

Data was analysed using MATLAB (R2016a, Mathworks, Massachusetts, USA).

## Results

### Administration of anaesthetics during early development has no gross effects on growth

Fig 2a displays worm lengths at the previously specified developmental stages for control and anaesthetic experiments. Results for anaesthetic experiments are pooled across the different drug concentrations. Boxplots give the distribution of lengths, with individual data points overlain. Coloured lines connect the median length at each developmental stage. Growth is as expected, and all animals follow similar trajectories. There is no significant difference in adult sizes (p>0.05 for every anaesthetic condition). Other size features, e.g. area, show a similar pattern (data not shown).

**Fig 2.**
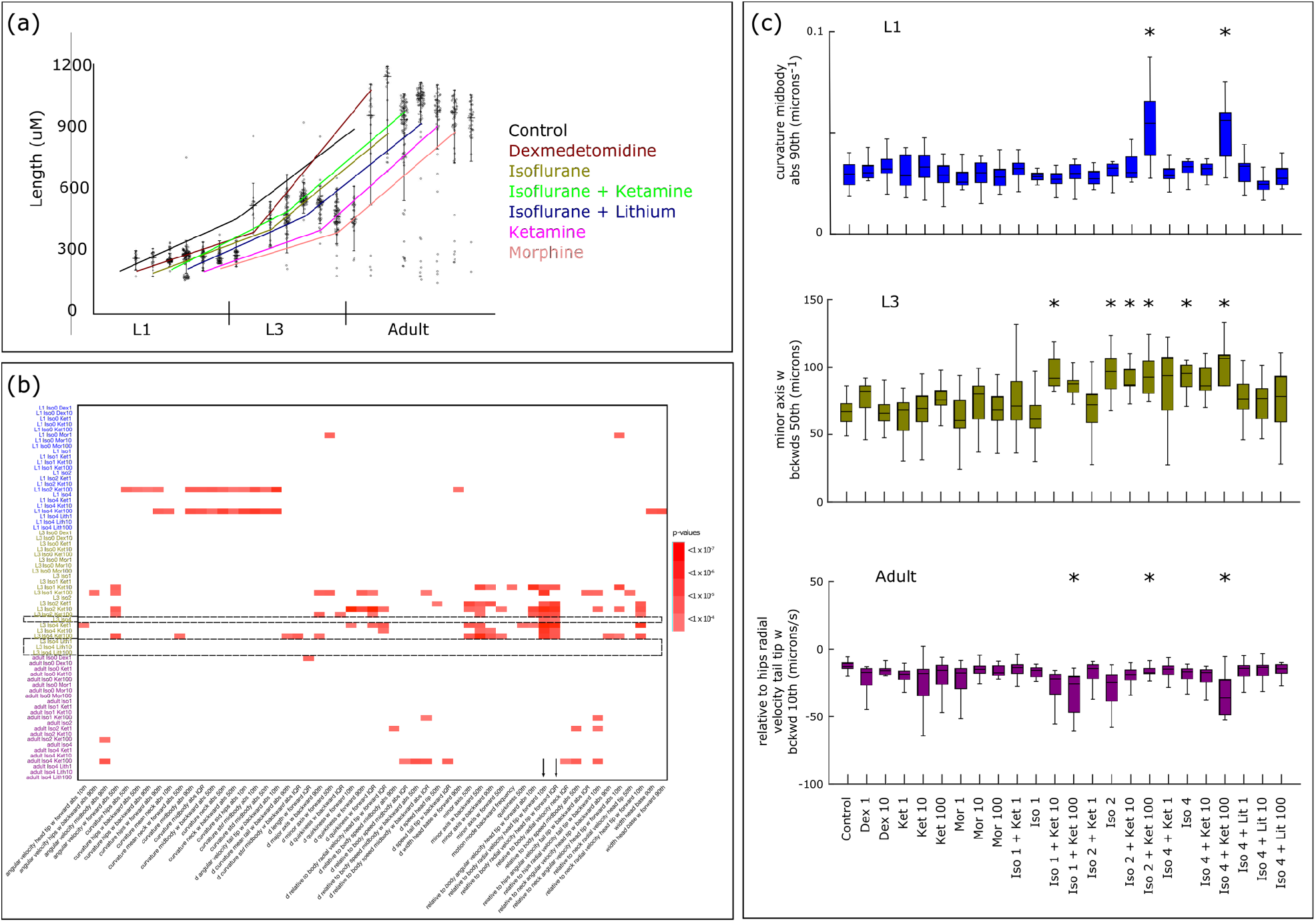
Early administration of general anaesthetics has no gross effects on growth but can have long term effects on behaviour. (a) Worm lengths at late L1, mid-L3, and mature adult stages. Results for different drug concentrations are pooled. Boxplots given distribution of lengths, with individual data points overlain. Coloured lines represent experimental condition, and connect median lengths at each developmental stage. Growth is as expected and all animals follow similar trajectories. There are no significant differences in adult size (p>0.05 for every condition). (b) Heatmap shows the p values of features that were found to be significantly different compared to control after correction for multiplicity, arranged by experimental condition and further divided by developmental stage. The behavioural phenotype of animals exposed to certain anaesthetic conditions varies from control, and this variation is developmental-stage-specific, i.e. changes in phenotype seen at one stage are not necessarily carried over to a later stage, and new changes can manifest as the animal develops. Dashed rectangles and arrows highlight two feature hits at L3 after exposure to Isoflurane; differences from control are abrogated after co-administration with lithium. (c) Examples of significant behaviour feature differences at L1 (top), L3 (middle) and adult (bottom). Dex = dexmedetomidine, Ket = ketamine, Mor = morphine, Iso = isoflurane; Lit = lithium. Numbers refer to either isoflurane concentration in percent air, or drug concentration in μM.

### There are developmental stage- and exposure-related variations in behaviour which persist long after emergence from anaesthesia

Fig 2b is a heat map of features that were found to be significantly different between anaesthesia-exposed animals and controls after correction for multiplicity. The data shows that the behavioural phenotype of animals exposed to certain anaesthetic conditions, but not others, varies from that of control animals. This variation is developmental-stage-specific, i.e. variations in phenotype seen at one stage of larval development are not necessarily carried over to a later stage, and new variations can manifest as the animal develops.

In total, there are 29 instances of a significant exposure/feature effect at L1, 81 at L3, and 15 at adult. Single-agent exposure with isoflurane yields 2 significant ‘hits’, both at L3 and at the highest 4% concentration only. There are no hits with single-agent exposure to ketamine at any concentration. However, the combination of isoflurane and ketamine leads to many persisting behavioural changes. There is evidence of a concentration effect – with exposure to higher concentrations causing more feature differences. The isoflurane 4% + ketamine 100μM combination leads to 13 feature hits at L1, 14 at L3, and 8 at adult. Other concentrations of the isoflurane + ketamine combination also show high numbers of feature hits. There are only 2 hits for morphine (at 1μM but not higher concentrations), and a single hit for dexmedetomidine (at 1μM but not higher) in total.

Fig 2c displays boxplots showing examples of significant feature differences at L1 (top panel), L3 (middle) and adult (bottom) stages. The latter findings show that behavioural changes persist after the period of neurogenesis, which is expected to be completed by L3.

### Co-administration of lithium rescues the behavioural effects of isoflurane administration

In this dataset, there are two feature hits resulting from exposure of larvae to Isoflurane 4%. Both are seen at L3 stage. In each case, co-administration of lithium at concentrations of 1, 10 or 100μM abrogated the differences with controls, without leading to any new difference in other features (Fig 2b and Fig 3).

**Fig 3.**
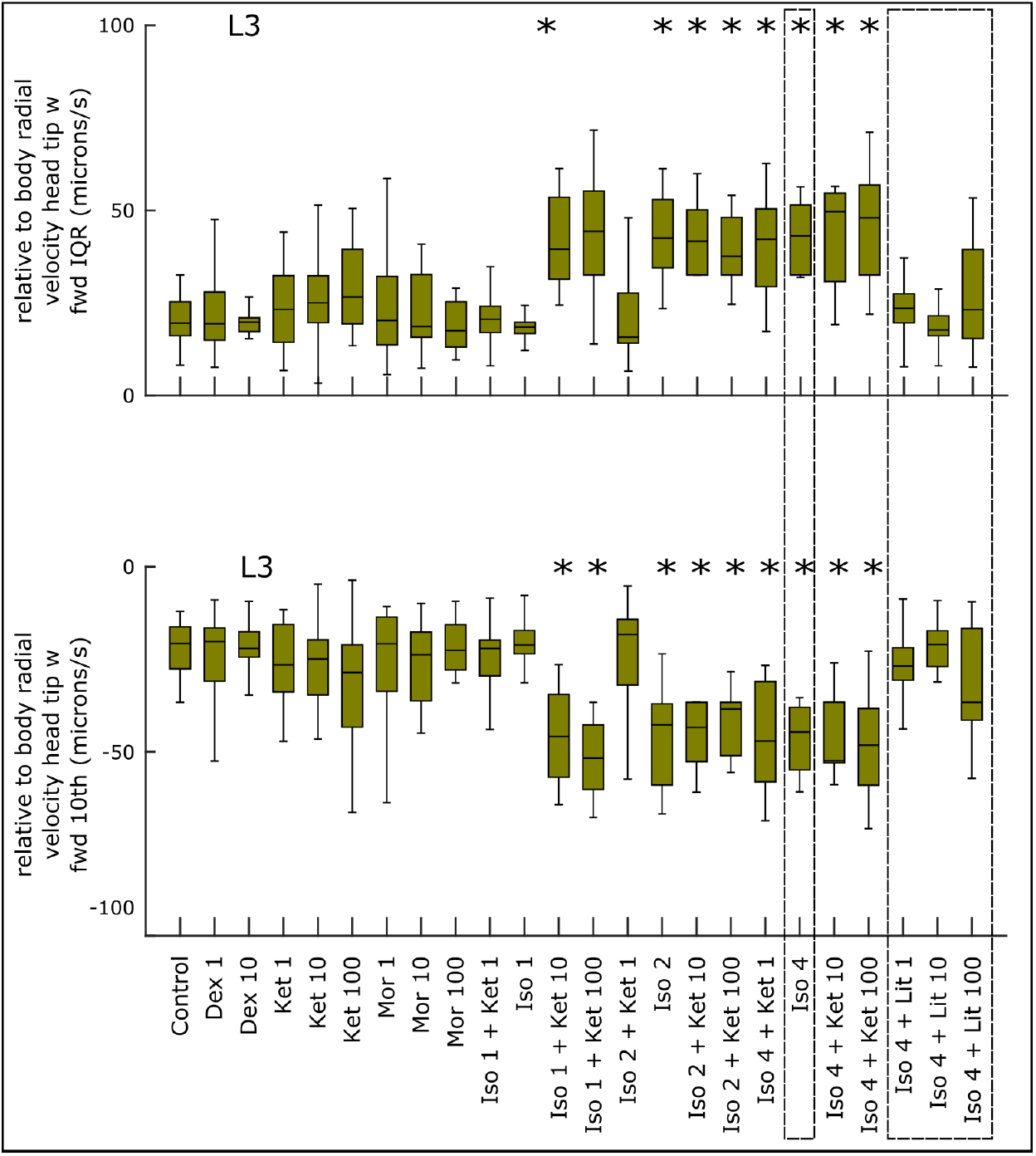
Boxplots of the two feature hits seen after administration of isoflurane alone. In both cases, coadministration of lithium abrogates the differences with controls.

## Discussion

This research establishes that *C. elegans* larvae exposed to anaesthesia shortly after hatching display later behavioural differences when compared to controls. The effect is agent- and concentrationdependent, with combinations of isoflurane and ketamine at higher doses consistently being correlated with significant differences. In this, the effect is similar to that seen in vertebrates, where the most robust and consistent findings of neurobehavioural impairment also come in experiments with multiple agents and stronger exposures^9,37,38^. We have also seen that morphine and dexmedetomidine, both agents which have been shown not to cause neuroapoptosis or behavioural change in vertebrates^31,32^, do not appear to cause lasting behavioural change in *C. elegans*. Although there are 2 hits for morphine and 1 for dexmedetomidine, the pattern of these, with hits in lower concentrations not being reproduced for higher concentrations, suggests that they are outliers.

There is some evidence here that lithium, which has been shown to rescue the neurotoxic phenotype in vertebrates^33^, may have a similar effect in *C. elegans*. Table 1 summarises the parallels between results from these experiments and vertebrate findings. The fact that results in the worm are broadly consistent with those in vertebrates suggests that these experiments could form the basis of a high-throughput screening system to detect compounds which could modulate anaesthesia-induced neurobehavioural impairment, and be used to identify mechanisms and genes involved in the phenomenon.

**Table 1.** Summary of evidence: anaesthetic exposure in C. elegans, and previous literature from vertebrate experiments.

| Anaesthetic conditions | C. elegans results | Vertebrate findings<sup>9,31,33,37,38,45–47</sup> |
| --- | --- | --- |
| Isoflurane | Evidence of behavioural impairment after exposure to 4% isoflurane as early L1, detectable at L3 (but not at late L1, and not persisting to adult) | Multiple studies from rodent and other animals, including non-human primates, provide evidence of histopathological changes and behavioural impairment. |
| Ketamine | Non-significant results in these experiments from single-agent exposure. | Evidence of histopathological change and behavioural impairment. |
| Morphine | No evidence of persisting behavioural change. | No evidence of histopathological or behavioural change. |
| Dexmedetomidine | No evidence of persisting behavioural change | No evidence of histopathological or behavioural change |
| Concentration effect | Isoflurane at 4% is associated with some behavioural feature changes at L3. Lower concentrations are not. | Higher dose-exposure parameters lead to increased histopathological and behavioural change. |
| Multi-drug cocktails | Ketamine/Isoflurane combination produces strongest evidence of persisting behavioural change. | Exposure to multi-drug cocktails (including combinations of volatile anaesthetics, ketamine, and benzodiazepines), lead to pronounced histopathological and behavioural change. |
| Lithium | Rescues behavioural effect seen with Isoflurane 4% | Evidence for rescuing isoflurane-induced neuroapoptosis in the infant primate brain. |

Many genes (around 80% by some estimates) and molecular pathways are known to be conserved between *C. elegans* and vertebrates^39,40^, and the worm has been used extensively for drug discovery and target identification. Forward genetic screens, which start with a phenotype of interest and aim to identify the genetic basis for the behaviour, have identified genes and products relevant to human disease phenotypes, including in neurodegenerative and neuromuscular disorders^41,42^. Chemistry-to-gene screens, in which mutagenized animals are exposed to a drug of interest and then screened for resistance to drug effect, have helped uncover clinically important molecular targets, such as those for antiparasitic drugs^43^. Gene-to-chemistry screens can facilitate the discovery of new drug targets by silencing genes in the search for resistance to a compound of interest. This approach led to the discovery that one compound in a library reduced muscular degeneration in a nematode model of Duchenne muscular dystrophy (DMD)^44^. The drug was the steroid prednisolone, in clinical use in DMD to palliate symptoms, and providing proof of principle of the potential of *C. elegans* to identify compounds for use in humans.

The advance of our mechanistic understanding of anaesthesia-induced neurobehavioural impairment, and the discovery of new compounds to ameliorate the phenomenon have both been recognised as research priorities. Existing methods have thus far produced an incomplete picture, and led to the identification of very few candidate drugs, most of which are years away from trials. It is possible to see how *C. elegans* could be used as an inexpensive, quick and tractable model for screening, with the aim of filtering out compounds or targets of interest for further validation. In contrast to cell line screening, whole-organism behavioural screens allow for unbiased investigation of a broad range of disease-related pathways, not just a cellular marker that is decided upon *a priori*. Given the gaps in our knowledge of the pathway towards behavioural impairment, this is an advantage.

Some approaches suggest themselves: the first is drug-discovery using libraries of pre-approved or novel compounds. These are unlikely to reveal new anaesthetic agents clearly, but may produce hits which enhance or supress the phenotype. Given the results seen in these experiments, a good starting point may be to expose worms as L1s to an isoflurane/ketamine/experimental drug combination, and measure for effect at L3, the stage at which the greatest perturbation of the phenotypic signature was recorded. We could speculate as to why L3 is stage at which behavioural impairment is most marked; perhaps the pertinent neural pathways are not fully developed by late-L1, and perhaps compensatory mechanisms have set in by adulthood despite the fact that neurogenesis is complete.

Further, mutagenesis of wildtype *C. elegans* may reveal mutants in whom the phenotype is enhanced or supressed, and provide a starting point for investigating molecular targets involved in the pathway. A complementary approach would be to employ a reverse genetic screen, using genome-wide RNAi libraries to systematically silence genes in the search for a mutant which is resistant to the compounds of interest – either the isoflurane/ketamine combination itself, or a drug which rescues the phenotype. With automated methods of dispensing worms and assay components, and new technologies for assay readouts and image analysis, such screens have significant potential to detect compounds and mechanisms of action which could have proven elusive to vertebrate research.

There are some limitations to this work. It is difficult to compare bioactivity in the nematode worm with human equivalents, because of the presence of the cuticle, uncertainty about pharmacokinetics and pharmacodynamics, and the effect of temperature on the potency of volatile anaesthetics. Only isoflurane/ketamine combinations show strong and persisting effects on behavioural change. This is broadly in keeping with vertebrate findings, although a few groups have demonstrated behavioural change with single-agent exposure alone (e.g. Lin et al^18^). It is possible that other *C. elegans* based screening protocols may be more sensitive to single-agent exposure, though this may not necessarily be an advantage if single-agent exposure does not lead to human impairment (as the best current evidence suggests^17^). In addition, although we have some evidence here that co-administration of lithium can abolish the behavioural effect seen with isoflurane exposure, it would be instructive to establish whether this rescue applies after treatment with combination exposures.

The chosen methods of phenotyping focus on features which are amenable to automated analysis, rather than high-order behaviours such as chemotaxis, which have previously been shown to be deficient after anaesthesia exposure^26^. However, this also allows unbiased screens, rather than making *a priori* judgements about which behaviours are likely to be mechanistically linked to the neuropathology induced by anaesthesia exposure.

This approach allows interrogation of phenotypic aspects of anaesthesia-induced neurobehavioural impairment, in the service of drug discovery and target identification. The use of quantitative phenotyping greatly enhances the power to detect ‘hits’ – compounds, genes and pathways of interest – over manual observation alone. These hits can then be further validated in vertebrate models or extended screens. Developments in robotics, microfluidics and image-analysis now allow for true high-throughput screening of nematodes^23,24,48^. *C. elegans* therefore is an excellent model with which to pursue drug discovery and mechanistic research. This work provides proof of principle for its use as a low-cost, high-throughput, whole-animal screening tool in the search for novel compounds and molecular mechanisms in AIN.

## Acknowledgements

This work was supported by the Medical Research Council through grant MC-A658-5TY30 to AEXB. We thank Dr Eleni Minga for help with analysis.

## Supplementary Data

**Table S1.**
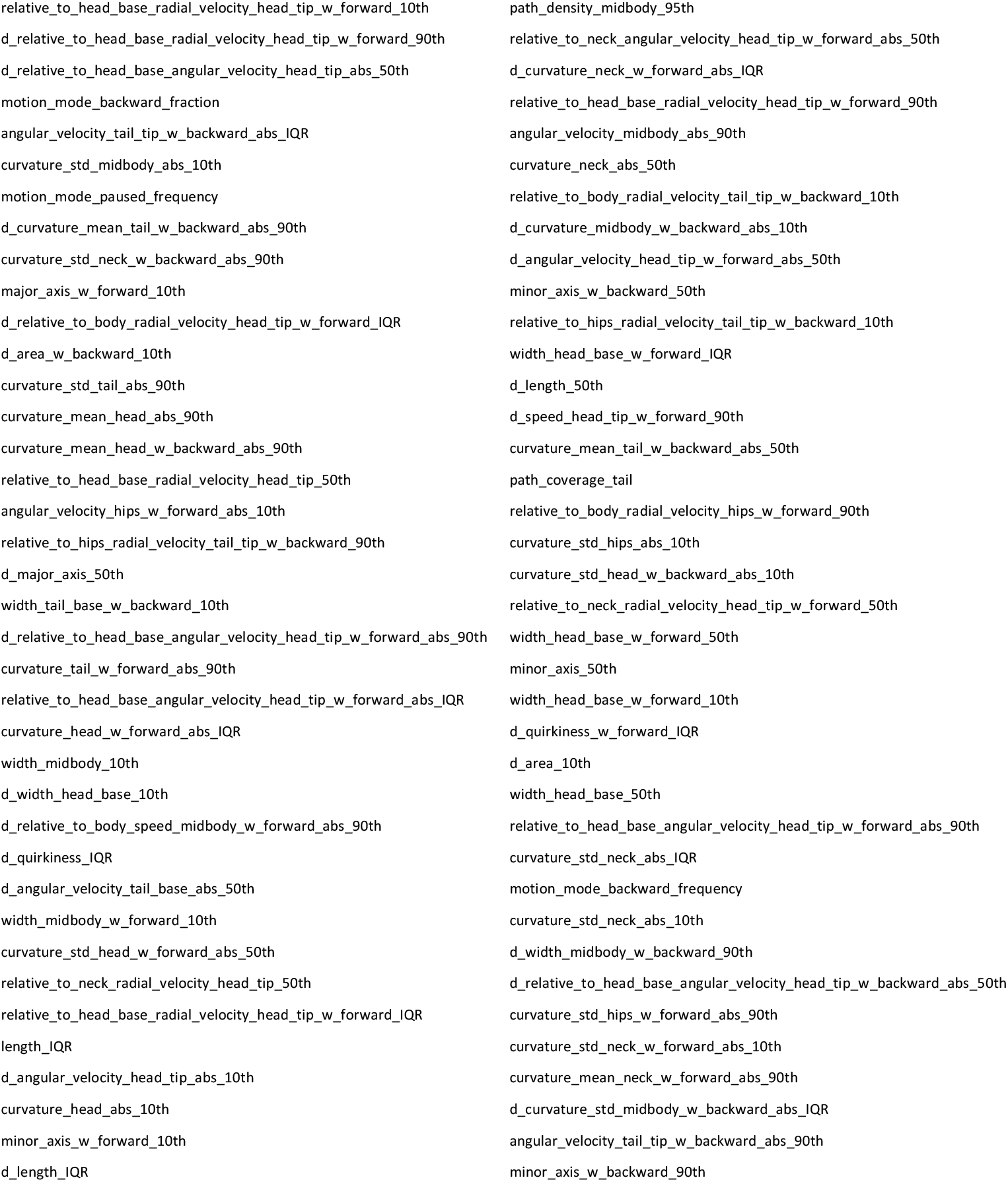
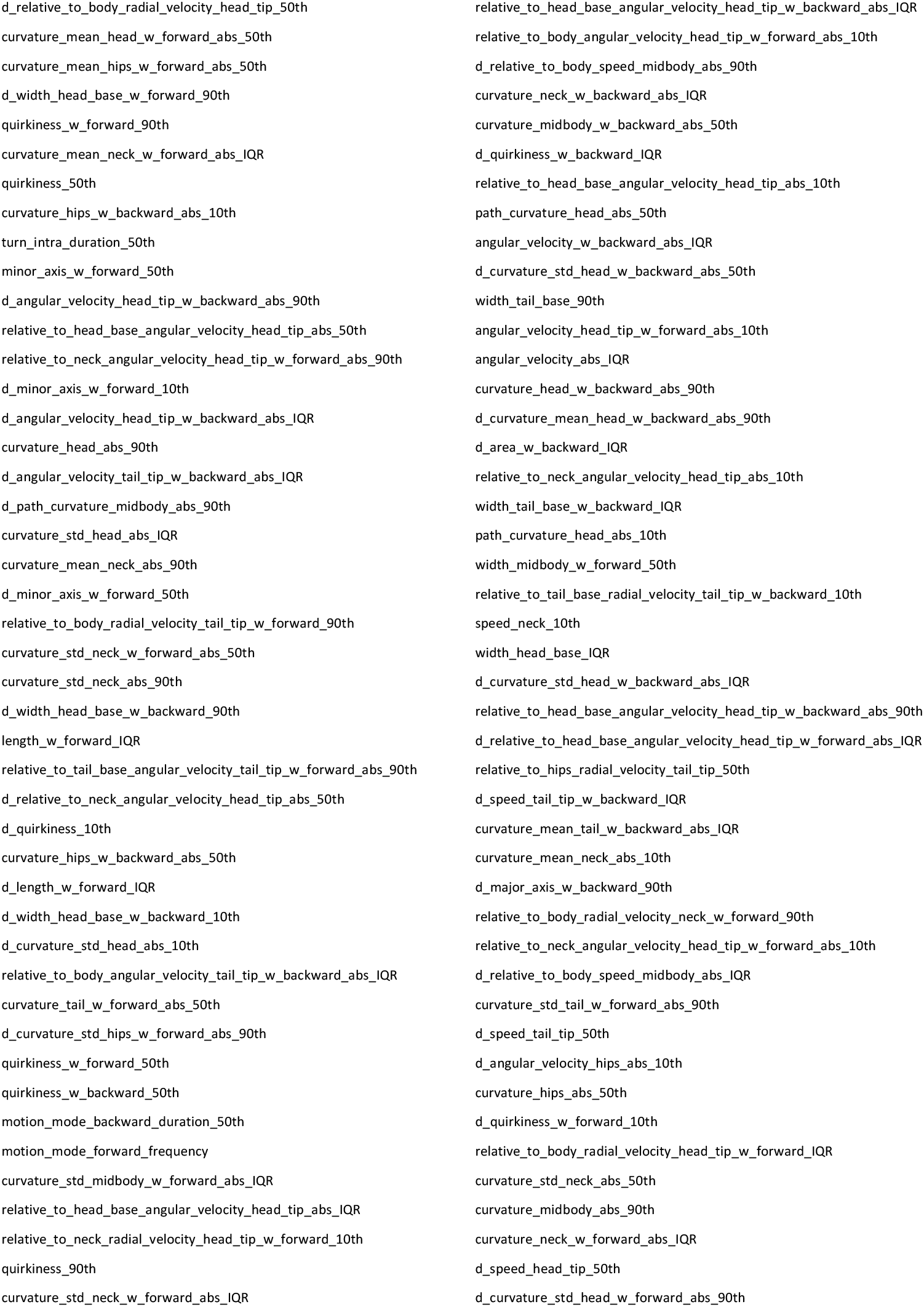
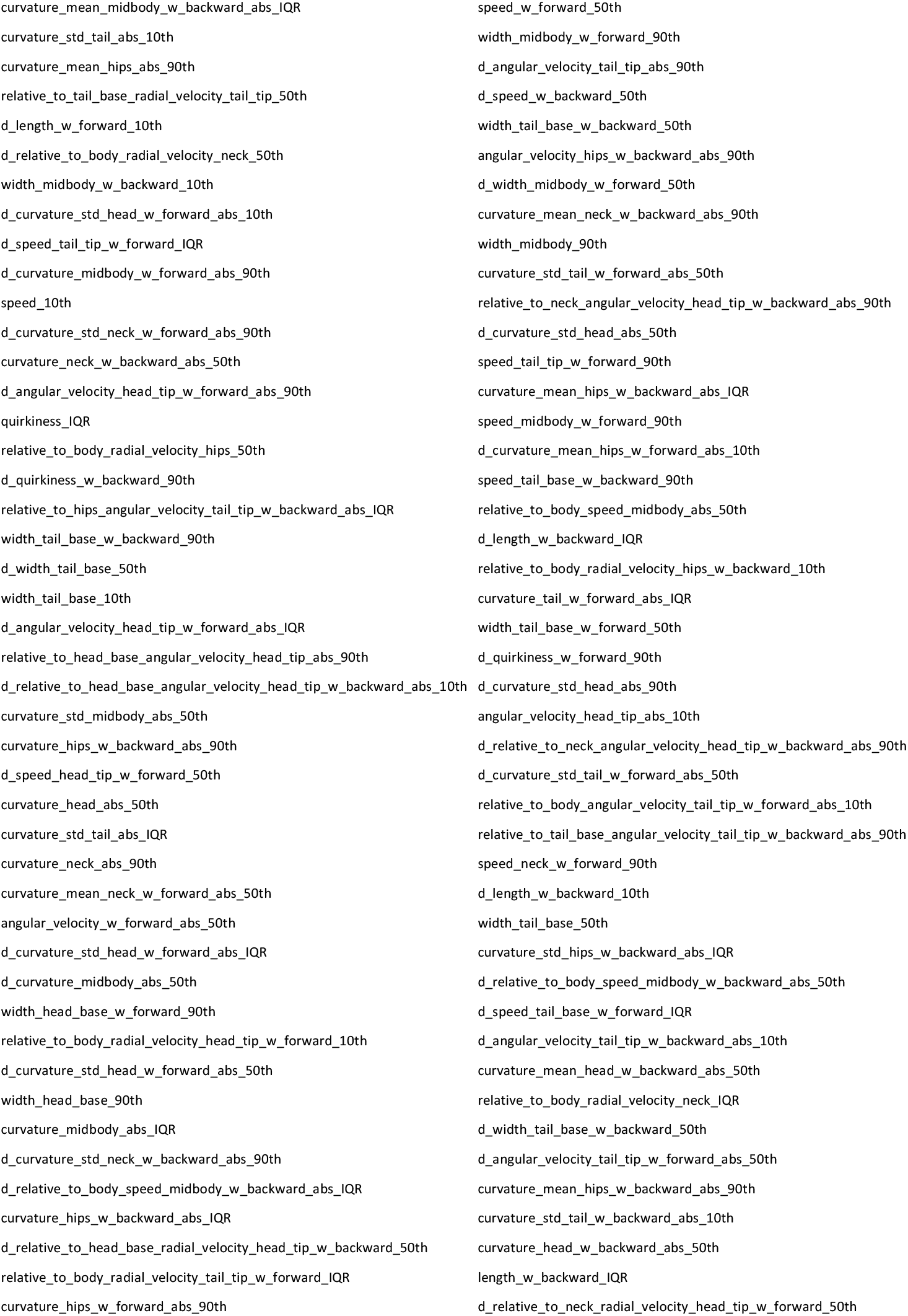
‘Tierpsy_256’ features^35^. There are many sets of 256 features which perform similarly in classification. The following is one such set, used in this analysis.

## References

1. Vutskits L. General Anesthesia: A Gateway to Modulate Synapse Formation and Neural Plasticity? Anesth Analg. 2012;115(5):1174–1182. doi:10.1213/ANE.0b013e31826a1178

2. Franks NP. General anaesthesia: from molecular targets to neuronal pathways of sleep and arousal. Nat Rev Neurosci. 2008;9(5):370–386. doi:10.1038/nrn2372

3. Bear MF, Kleinschmidt A, Gu QA, Singer W. Disruption of experience-dependent synaptic modifications in striate cortex by infusion of an NMDA receptor antagonist. J Neurosci. 1990;10(3):909–925. doi:10.1523/JNEUROSCI.10-03-00909.1990

4. Hensch TK, Fagiolini M, Mataga N, Stryker MP, Baekkeskov S, Kash SF. Local GABA Circuit Control of Experience-Dependent Plasticity in Developing Visual Cortex. Science. 1998;282(5393):1504–1508. doi:10.1126/science.282.5393.1504

5. Malenka RC, Bear MF. LTP and LTD: An Embarrassment of Riches. Neuron. 2004;44(1):5–21. doi:10.1016/j.neuron.2004.09.012

6. Lener MS, Kadriu B, Zarate CA. Ketamine and Beyond: Investigations into the Potential of Glutamatergic Agents to Treat Depression. Drugs. 2017;77(4):381–401. doi:10.1007/s40265-017-0702-8

7. Andropoulos DB. Anesthesia and Developing Brains — Implications of the FDA Warning. N Engl J Med. 2017;376:905–907. doi:https://doi.org/10.1056/NEJMp1700196

8. Ikonomidou C, Bosch F, Miksa M, et al. Blockade of NMDA Receptors and Apoptotic Neurodegeneration in the Developing Brain. Science. 1999;283(5398):70–74. doi:10.1126/science.283.5398.70

9. Jevtovic-Todorovic V, Hartman RE, Izumi Y, et al. Early Exposure to Common Anesthetic Agents Causes Widespread Neurodegeneration in the Developing Rat Brain and Persistent Learning Deficits. J Neurosci. 2003;23(3):876–882. doi:10.1523/JNEUROSCI.23-03-00876.2003

10. Cattano D, Young C, Straiko MMW, Olney JW. Subanesthetic doses of propofol induce neuroapoptosis in the infant mouse brain. Anesth Analg. 2008;106(6):1712–1714. doi:10.1213/ane.0b013e318172ba0a

11. Paule MG, Li M, Allen RR, et al. Ketamine anesthesia during the first week of life can cause long-lasting cognitive deficits in rhesus monkeys. Neurotoxicol Teratol. 2011;33(2):220–230. doi:10.1016/j.ntt.2011.01.001

12. Walkden G, Pickering A, Gill H. Assessing Long-term Neurodevelopmental Outcome Following General Anesthesia in Early Childhood: Challenges and Opportunities. Anesth Analg. 2019;128(4):681–694. doi:10.1213/ANE.0000000000004052

13. DiMaggio C, Sun LS, Kakavouli A, Byrne MW, Li G. A Retrospective Cohort Study of the Association of Anesthesia and Hernia Repair Surgery With Behavioral and Developmental Disorders in Young Children. J Neurosurg Anesthesiol. 2009;21(4):286–291. doi:10.1097/ANA.0b013e3181a71f11

14. Wilder RT, Flick RP, Sprung J, et al. Early Exposure to Anesthesia and Learning Disabilities in a Population-based Birth Cohort. Anesthesiology. 2009;110(4):796–804. doi:10.1097/01.anes.0000344728.34332.5d

15. DiMaggio C, Sun LS, Ing C, Li G. Pediatric anesthesia and neurodevelopmental impairments: a Bayesian meta-analysis. J Neurosurg Anesthesiol. 2012;24(4):376–381. doi:10.1097/ANA.0b013e31826a038d

16. Sun LS, Li G, Miller TLK, et al. Association Between a Single General Anesthesia Exposure Before Age 36 Months and Neurocognitive Outcomes in Later Childhood. JAMA. 2016;315(21):2312. doi:10.1001/jama.2016.6967

17. McCann ME, Graaff JC de, Dorris L, et al. Neurodevelopmental outcome at 5 years of age after general anaesthesia or awake-regional anaesthesia in infancy (GAS): an international, multicentre, randomised, controlled equivalence trial. The Lancet. 2019;393(10172):664–677. doi:10.1016/S0140-6736(18)32485-1

18. Lin D, Liu J, Kramberg L, Ruggiero A, Cottrell J, Kass IS. Early-life single-episode sevoflurane exposure impairs social behavior and cognition later in life. Brain Behav. 2016;6(9):e00514. doi:10.1002/brb3.514

19. Ji M, Wang X, Sun X, et al. Environmental Enrichment Ameliorates Neonatal Sevoflurane Exposure-Induced Cognitive and Synaptic Plasticity Impairments. J Mol Neurosci. 2015;57(3):358–365. doi:10.1007/s12031-015-0627-1

20. Li J, Xiong M, Nadavaluru PR, et al. Dexmedetomidine Attenuates Neurotoxicity Induced by Prenatal Propofol Exposure. J Neurosurg Anesthesiol. 2015;28:51–64). doi:10.1097/ANA.0000000000000181

21. Zheng H, Dong Y, Xu Z, et al. Sevoflurane Anesthesia in Pregnant Mice Induces Neurotoxicity in Fetal and Offspring Mice. Anesthesiology. 2013;118(3):516–526. doi:10.1097/ALN.0b013e3182834d5d

22. Zhao Y, Chen K, Shen X. Environmental Enrichment Attenuated Sevoflurane-Induced Neurotoxicity through the PPAR-γ Signaling Pathway. BioMed Research International. doi:https://doi.org/10.1155/2015/107149

23. Cornaglia M, Lehnert T, Gijs MAM. Microfluidic systems for high-throughput and high-content screening using the nematode Caenorhabditis elegans. Lab Chip. 2017;17(22):3736–3759. doi:10.1039/C7LC00509A

24. de Carlos Cáceres I, Porto DA, Gallotta I, et al. Automated screening of C. elegans neurodegeneration mutants enabled by microfluidics and image analysis algorithms. Integr Biol. 2018;10(9):539–548. doi:10.1039/c8ib00091c

25. Conradt B. Programmed cell death. WormBook. Published online 2005. doi:10.1895/wormbook.1.32.1

26. Gentry KR, Steele LM, Sedensky MM, Morgan PG. Early Developmental Exposure to Volatile Anesthetics Causes Behavioral Defects in Caenorhabditis elegans. Anesth Analg. 2013;116(1):185–189. doi:10.1213/ANE.0b013e31826d37c5

27. Yu D, Li L, Yuan W. Neonatal anesthetic neurotoxicity: Insight into the molecular mechanisms of long-term neurocognitive deficits. Biomed Pharmacother. 2017;87:196–199. doi:10.1016/j.biopha.2016.12.062

28. Johnson SC, Pan A, Li L, Sedensky M, Morgan P. Neurotoxicity of anesthetics: Mechanisms and meaning from mouse intervention studies. Neurotoxicol Teratol. 2019;71:22–31. doi:10.1016/j.ntt.2018.11.004

29. Stiernagle T. Maintenance of C. elegans. WormBook. Published online 2006. doi:10.1895/wormbook.1.101.1

30. Islam P. Making OP50 solution from Frozen Stock. Published online November 23, 2018. doi:10.17504/protocols.io.vube6sn

31. Schuurmans J, Benders M, Lemmers P, Bel F van. Neonatal morphine in extremely and very preterm neonates: its effect on the developing brain – a review. J Matern Fetal Neonatal Med. 2015;28(2):222–228. doi:10.3109/14767058.2014.908178

32. van Hoorn CE, Hoeks SE, Essink H, Tibboel D, de Graaff JC. A systematic review and narrative synthesis on the histological and neurobehavioral long-term effects of dexmedetomidine. Paediatr Anaesth. 2019;29(2):125–136. doi:10.1111/pan.13553

33. Noguchi KK, Johnson SA, Kristich LE, et al. Lithium Protects Against Anaesthesia Neurotoxicity In The Infant Primate Brain. Sci Rep. 2016;6(1):22427. doi:10.1038/srep22427

34. Javer A, Currie M, Lee CW, et al. An open-source platform for analyzing and sharing worm-behavior data. Nat Methods. 2018;15(9):645–646. doi:10.1038/s41592-018-0112-1

35. Javer A, Ripoll-Sánchez L, Brown AEX. Powerful and interpretable behavioural features for quantitative phenotyping of Caenorhabditis elegans. Philos Trans R Soc B Biol Sci. 2018;373(1758):20170375. doi:10.1098/rstb.2017.0375

36. Benjamini Y, Hochberg Y. Controlling the False Discovery Rate: A Practical and Powerful Approach to Multiple Testing. J R Stat Soc Ser B Methodol. 1995;57(1):289–300.

37. Yon J-H, Daniel-Johnson J, Carter LB, Jevtovic-Todorovic V. Anesthesia induces neuronal cell death in the developing rat brain via the intrinsic and extrinsic apoptotic pathways. Neuroscience. 2005;135(3):815–827. doi:10.1016/j.neuroscience.2005.03.064

38. Lunardi N, Ori C, Erisir A, Jevtovic-Todorovic V. General Anesthesia Causes Long-Lasting Disturbances in the Ultrastructural Properties of Developing Synapses in Young Rats. Neurotox Res. 2010;17(2):179–188. doi:10.1007/s12640-009-9088-z

39. Lai C-H, Chou C-Y, Ch’ang L-Y, Liu C-S, Lin W. Identification of Novel Human Genes Evolutionarily Conserved in Caenorhabditis elegans by Comparative Proteomics. Genome Res. 2000;10(5):703–713. doi:10.1101/gr.10.5.703

40. Kim W, Underwood RS, Greenwald I, Shaye DD. OrthoList 2: A New Comparative Genomic Analysis of Human and Caenorhabditis elegans Genes. Genetics. 2018;210(2):445–461. doi:10.1534/genetics.118.301307

41. Dimitriadi M, Hart AC. Neurodegenerative disorders: Insights from the nematode Caenorhabditis elegans. Neurobiol Dis. 2010;40(1):4–11. doi:10.1016/j.nbd.2010.05.012

42. Sleigh J, Sattelle D. C. elegans models of neuromuscular diseases expedite translational research. Transl Neurosci. 2010;1(3):214–227. doi:10.2478/v10134-010-0032-9

43. Jones AK, Buckingham SD, Sattelle DB. Chemistry-to-gene screens in Caenorhabditis elegans. Nat Rev Drug Discov. 2005;4(4):321–330. doi:10.1038/nrd1692

44. Gaud A, Simon J-M, Witzel T, Carre-Pierrat M, Wermuth CG, Ségalat L. Prednisone reduces muscle degeneration in dystrophin-deficient Caenorhabditis elegans. Neuromuscul Disord. 2004;14(6):365–370. doi:10.1016/j.nmd.2004.02.011

45. Zou X, Liu F, Zhang X, et al. Inhalation anesthetic-induced neuronal damage in the developing rhesus monkey. Neurotoxicol Teratol. 2011;33(5):592–597. doi:10.1016/j.ntt.2011.06.003

46. Shen X, Dong Y, Xu Z, et al. Selective Anesthesia-induced Neuroinflammation in Developing Mouse Brain and Cognitive Impairment. Anesthesiology. 2013;118(3):502–515. doi:10.1097/ALN.0b013e3182834d77

47. Boscolo A, Milanovic D, Starr J, et al. Early Exposure to General Anesthesia Disturbs Mitochondrial Fission and Fusion in the Developing Rat Brain. Anesthesiology. 2013;118(5):1086–1097. doi:10.1097/ALN.0b013e318289bc9b

48. Rajamuthiah R, Fuchs BB, Jayamani E, et al. Whole Animal Automated Platform for Drug Discovery against Multi-Drug Resistant Staphylococcus aureus. PLOS ONE. 2014;9(2):e89189. doi:10.1371/journal.pone.0089189

